# Label-free 3D subcellular phenotyping of mouse embryos by holotomography enables early prediction of blastocyst formation

**DOI:** 10.1101/2024.05.07.592317

**Authors:** Chungha Lee, Geon Kim, Taeseop Shin, Sangho Lee, Jae Young Kim, Kyoung Hee Choi, Jieun Do, Jaehyeong Park, Jaephil Do, Ji Hyang Kim, YongKeun Park

## Abstract

Accurate embryo quality assessment is central to improving outcomes in in vitro fertilization (IVF), yet current practice relies mainly on subjective two-dimensional (2D) morphology. Here we present a label-free framework for quantitative three-dimensional (3D) embryo phenotyping using low-coherence holotomography (HT). Time-lapse HT enabled volumetric imaging of mouse embryos from the 2-cell stage to the blastocyst without affecting developmental competence, capturing subcellular features at high resolution. Quantitative analysis revealed that matured embryos exhibited higher blastomere counts, greater spatial variability, and tighter nuclear packing, whereas arrested embryos showed enlarged blastomeres, elevated cytoplasmic heterogeneity, and fewer, larger nuclei. Machine learning models trained on these features achieved robust prediction of blastocyst formation (AUC up to 0.958). Together, these findings demonstrate that HT provides objective and interpretable 3D biomarkers that could augment and transform embryo selection in IVF.

## Introduction

Despite the widespread use of assisted reproductive technology (ART), live birth rates per in vitro fertilization (IVF) cycle remain modest at ∼30% (*1*). A key determinant of success is accurate embryo quality assessment, yet current clinical practice relies largely on morphological grading of blastocysts on day 5 (*2, 3*). Time-lapse imaging has introduced dynamic parameters such as cleavage timing and division synchrony (*4, 5*), but these approaches remain constrained to two-dimensional (2D) observations. Because embryos are inherently three-dimensional (3D) and structurally complex, 2D profiling provides only limited insight into their true developmental potential.

Fluorescence-based 3D microscopy can reveal subcellular detail (*6*–*11*) but is incompatible with clinical IVF due to labeling requirements and concerns over phototoxicity. Consequently, label-free approaches such as bright-field, dark-field, and Hoffman modulation contrast microscopy are widely adopted in practice (*4, 12*), yet these modalities remain qualitative and limited to surface-level features. This gap highlights the need for quantitative, label-free methods capable of capturing embryo architecture in 3D at subcellular resolution. Recent advances in stem-cell-derived embryo models have further enabled high-resolution visualization of peri-implantation and gastrulation-like stages *in vitro* (*13, 14*), accentuating the need for label-free approaches that can capture comparable structural detail without genetic perturbation.

Quantitative phase imaging (QPI) provides such a capability by exploiting the refractive index (RI) as an intrinsic source of contrast (*15*–*17*). Holotomography (HT), a 3D implementation of QPI (*18*), has proven effective for imaging 3D biology specimens, including bacteria (*19, 20*), cells (*21*–*25*), organoids (*26*–*28*), tissues (*29*–*33*), and embryos across species (*34*–*40*). While previous studies have demonstrated the feasibility of HT for embryo imaging, the 3D subcellular dynamics that underlie developmental competence remain largely unexplored.

In parallel, artificial intelligence (AI) has emerged as a powerful tool for embryo selection (*1*). Most AI-based models rely on morphokinetic annotations from time-lapse imaging (*5, 41*) or static day-5 morphology (*42, 43*), and have been trained using implantation data (*44, 45*), maternal age (*46*), or preimplantation genetic testing results (*46, 47*). These models, however, primarily emphasize cell-division timing and gross 2D morphology, with limited integration of quantitative subcellular features.

Here, we present a systematic framework for quantitative, label-free, high-resolution 3D imaging of preimplantation embryos. Using low-coherence HT, we acquired RI tomograms of mouse embryos from the 2-cell stage to the expanded blastocyst and correlated early subcellular configurations with developmental outcomes. We identify quantitative parameters describing blastomere organization, nuclear architecture, and cytoplasmic heterogeneity that distinguish embryos with divergent fates. Leveraging these features, we establish predictive models of blastocyst formation, highlighting the potential of 3D subcellular phenotyping as a powerful noninvasive biomarker for embryo quality.

## Results

### Time-lapse HT of mouse preimplantation embryos

To investigate early embryonic development with quantitative, label-free imaging, we implemented a customized low-coherence HT system optimized for preimplantation embryos (Fig. 1A). The system employed patterned red-light illumination (624 nm) projected via a digital micromirror device and condenser lens, followed by axial scanning and 3D deconvolution to reconstruct RI tomograms (See Methods). Embryos were cultured in a dedicated incubation chamber, and groups of embryos were imaged over a 72-hour period from the 2-cell stage through the expanded blastocyst stage. The reconstructed tomograms provided volumetric information with subcellular resolution, enabling quantitative analysis of dynamic developmental features (Figs. 1B–C).

**Fig. 1:**
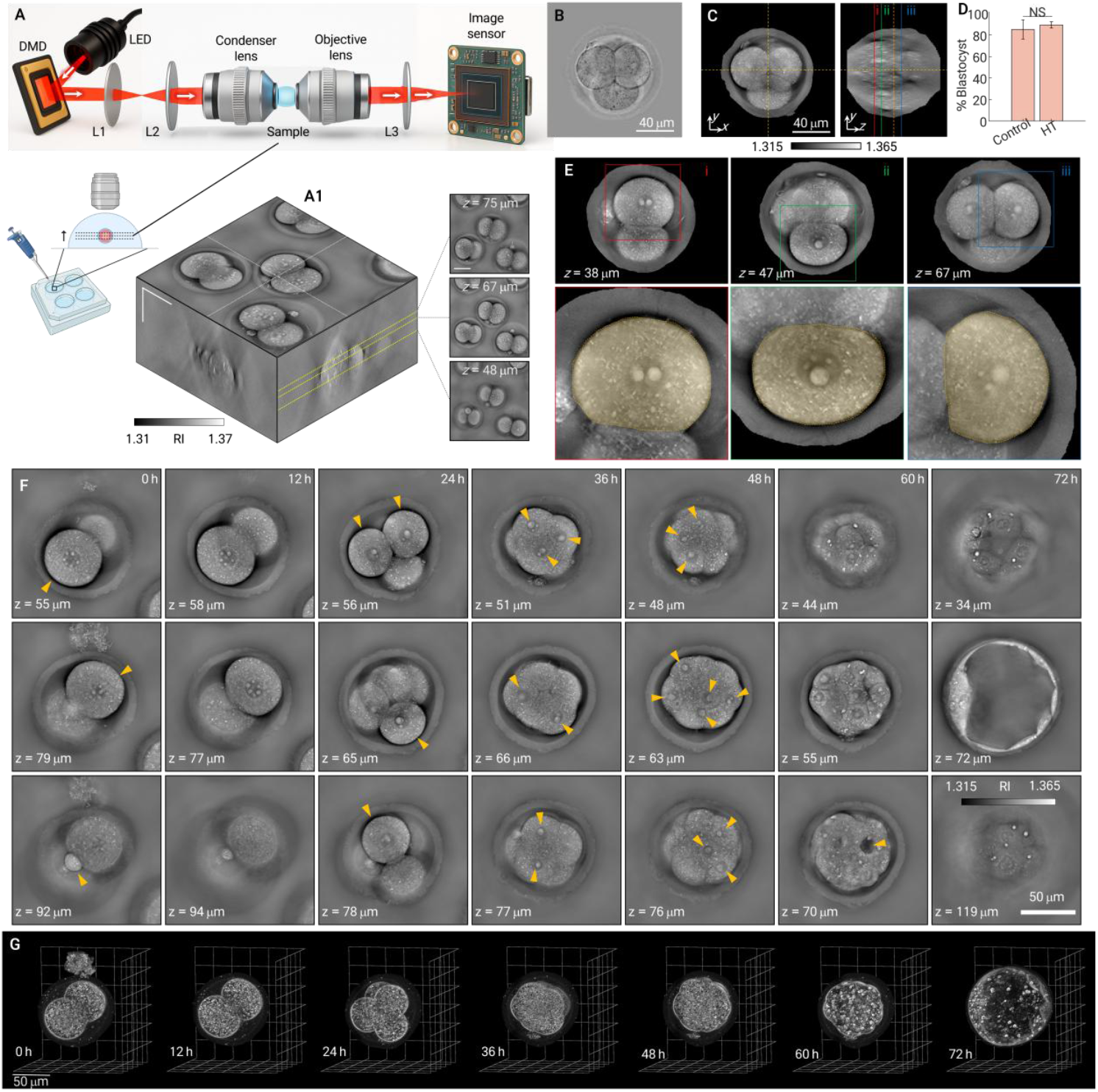
Time-lapse HT of mouse preimplantation embryos. **A**, Overview of the imaging system and workflow. The illumination pattern from a 624-nm LED is controlled using a DMD and relayed to the condenser back focal plane to generate four optimized illumination patterns; transmitted light is collected by the objective and recorded on a CMOS sensor. Embryos are cultured in group culture within a 4-well dish mounted on a heated/CO2-controlled stage (inset, **A1**). Representative reconstructed 3D RI tomogram showing the imaging volume and axial sections used for quantitative analysis. **B**, Bright-field reference image at the 2-cell stage. **C**, Orthogonal RI sections illustrating optical sectioning and subcellular contrast in HT (left, x–y; middle, x–z). **D**, comparison of blastocyst formation rates for control vs HT-imaged embryos showing no significant difference (NS), confirming assay safety. **E**, HT cross-sections of blastomeres at different focal planes (i: *z* = 38 μm, ii: *z* = 47 μm, iii: *z* = 67 μm), with cytoplasmic boundaries outlined (yellow dashed lines). Insets highlight corresponding planes in the embryo. **F**, Representative HT x–y slices acquired every 12 h from 0 to 72 h post 2-cell stage (see Movie S1). Yellow arrowheads mark key developmental features resolved in 3D: two blastomeres and polar body at distinct focal planes (*t* = 0 h, yellow arrows), multiple blastomeres at their respective planes (*t* = 24 h, yellow arrow), nuclei during compaction (*t* = 36 h and *t* = 48 h, yellow arrows), cavity formation (*t* = 60 h, yellow arrows), and trophectoderm with inner cell mass in the expanded blastocyst (*t* = 72 h). *z* denotes the focal depth of each slice. Scale bar (last panel), 50 µm. **G**, Gradient-based 3D renderings of the same embryo across time, highlighting evolving cellular architecture from 2-cell to expanded blastocyst.

We first assessed the safety and feasibility of HT for continuous embryo monitoring. Blastocyst formation rates were compared between embryos cultured under conventional conditions and those subjected to HT imaging. No significant differences were observed (Control: 85.15 ± 2.92%; HT: 89.34 ± 8.92%; Fig. 1D), indicating that repeated exposure to red light in the system did not impair developmental competence. Both values meet the FDA guidelines for mouse embryo assays in ART device testing (*48*), supporting the safety and feasibility of our approach. This result is consistent with prior reports of red-light safety in mammalian embryos and validates HT for long-term, noninvasive imaging of embryo development.

To further examine the spatial organization of blastomeres, we analyzed RI tomograms at multiple focal depths (*z* = 38, 47, and 67 μm) (Fig. 1E). Distinct blastomeres were clearly resolved at each plane, enabling direct visualization of both nuclear and cytoplasmic structures in three dimensions. The volumetric HT images delineated the cytoplasmic boundaries of individual blastomeres, which were outlined for quantitative assessment. This 3D information allowed reliable segmentation and measurement of blastomere size and morphology, highlighting the ability of HT to extract quantitative parameters beyond conventional 2D observation.

Representative RI tomograms captured key developmental milestones across the 72-hour culture period (Figs. 1F–G). At the 2-cell stage, distinct blastomeres and polar bodies were clearly resolved in different focal planes, supporting accurate volumetric measurement of individual cells. As embryos progressed to the 4- and 8-cell stages, HT provided detailed visualization of blastomere morphology, including nuclei and nucleoli during compaction. By 60 h, the onset of cavity formation was evident, and by 72 h, expanded blastocysts were imaged with sufficient contrast to reveal the inner cell mass and trophectoderm. The ability to visualize these structures in three dimensions highlights the power of HT to capture dynamic subcellular changes that are inaccessible to conventional 2D bright-field microscopy.

Finally, gradient-based renderings of the reconstructed RI tomograms further emphasized the unique capability of HT to provide both global morphology and local subcellular detail (Fig. 1G). Unlike 2D imaging, which compresses spatial information into a single plane, HT captured the volumetric organization of blastomeres, nuclei, and emerging blastocoel cavities throughout development. These observations establish HT as a robust platform for time-lapse, label-free imaging of preimplantation embryos, laying the foundation for quantitative analysis of developmental trajectories in subsequent experiments.

### 3D visualization of subcellular features distinguishing matured and arrested embryos

To investigate structural differences associated with developmental competence, we compared RI tomograms of embryos that successfully developed to expanded blastocysts within 72 h (matured group) and embryos that failed to reach the blastocyst stage (arrested group) (Fig. 2). Both bright-field images and HT-derived 3D reconstructions were analyzed, allowing simultaneous evaluation of classical morphological markers and subcellular features that are inaccessible by conventional microscopy. This approach enabled systematic comparison across cleavage, compaction, and blastocyst formation stages.

**Fig. 2:**
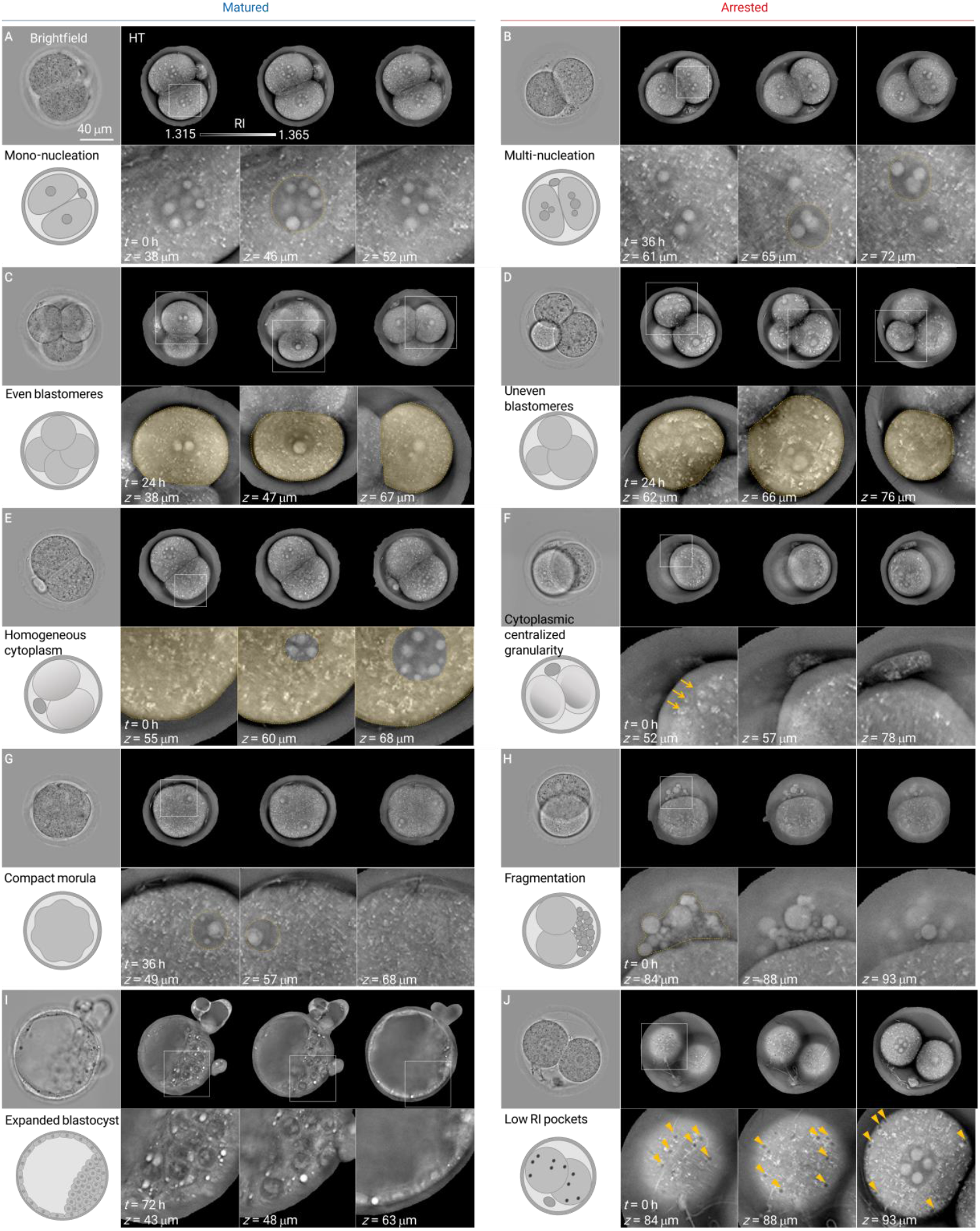
3D visualization of subcellular features in embryos with different developmental outcomes. Representative bright-field images (left) and HT images (right) of the matured embryos (blue, panels **A, C, E, G, I**) that developed to expanded blastocysts within 72 h, and the arrested embryos (red, panels **B, D, F, H, J**) that failed to reach the blastocyst stage. **A–B**, Detection of mono-nucleation (**A**) versus multi-nucleation (**B**, yellow dashed outlines) in 2-cell embryos. **C–D**, Blastomere morphology showing uniform size (**C**) compared to uneven blastomere size (**D**). **E–F**, Cytoplasmic characteristics, with homogeneous cytoplasm in matured embryos (**E**) versus irregular cytoplasmic granularity in arrested embryos (**F**, yellow arrowheads). **G–H**, Nuclear visualization in compacted morulae (**G**) and fragmentation in arrested embryos (**H**). **I–J**, Expanded blastocyst structures including blastocoel cavity, inner cell mass, and trophectoderm (**I**), and dispersed low-RI pockets in the cytoplasm of arrested embryos (**J**, yellow arrowheads). Each panel shows bright-field reference images (top left), schematic diagrams (bottom left), and corresponding HT slices or magnified RI sections (right).

Multi-nucleation and blastomere size variation are established morphological indicators of embryo quality (*3, 12*). The present approach enabled these features to be examined with greater clarity in three dimensions. At the 2-cell stage, HT revealed distinct patterns of nuclear morphology between groups. Matured embryos exhibited mono-nucleation, consistent with normal cleavage divisions (Fig. 2A), whereas multi-nucleation was observed exclusively in arrested embryos (Fig. 2B), a well-established marker of reduced developmental potential. The optical sectioning capability of HT facilitated reliable detection of these abnormalities across focal planes, avoiding ambiguity from 2D projections.

Analysis of blastomere morphology further highlighted differences between groups. In matured embryos, blastomeres displayed uniform size and shape (Fig. 2C), whereas increased variability in blastomere size was sporadically observed in arrested embryos (Fig. 2D). These observations align with consensus grading criteria that identify uneven blastomere size as a negative prognostic indicator. Importantly, HT allowed quantitative delineation of blastomere boundaries, suggesting that volumetric size variation could be systematically measured rather than subjectively judged.

Cytoplasmic texture provided another discriminating feature. During early cleavage stages, matured embryos generally exhibited homogeneous cytoplasm with smooth RI distributions (Fig. 2E). In contrast, arrested embryos often exhibited cytoplasmic irregularities such as localized or centrally concentrated granularity (Fig. 2F, yellow arrowheads). Although conventional guidelines caution against overinterpretation of cytoplasmic morphology, our HT-based 3D visualizations demonstrate that heterogeneity can be objectively assessed and may contribute additional predictive value.

HT enabled visualization of complex subcellular and multicellular structures during later stages of development. In matured embryos, morulae exhibited compact cellular organization with well-aligned nuclei (Fig. 2G), and expanded blastocysts displayed clear formation of the blastocoel cavity, inner cell mass (ICM), and trophectoderm (TE) (Fig. 2I). The 3D RI tomograms provided volumetric context for cavity expansion and ICM–TE relationships, illustrating how HT captures both classical morphological markers and additional structural details relevant to embryo quality.

Compared to matured embryos, arrested embryos showed irregular features in a subset of embryos, including fragmentation (Fig. 2H) and dispersed low-RI pockets within the cytoplasm (Fig. 2J, yellow arrowheads). These abnormalities, often unresolved in bright-field microscopy, were clearly identified by HT and likely reflect cellular stress underlying developmental arrest. Such low-RI cytoplasmic pockets may represent local fluid compartments or vacuolation associated with dysregulated hydraulic balance, a mechanism increasingly recognized as central to embryonic and reproductive tissue morphogenesis (*49*). Together, these results emphasize the ability of HT to reveal both recognized and novel subcellular abnormalities for comprehensive embryo assessment.

### Noninvasive 3D subcellular assessment at 24 h post–2-cell stage

The developmental trajectories of embryos with different outcomes became distinct at 24 h post–2-cell stage. We therefore focused on this time point to perform a detailed analysis of physiologically relevant 3D morpho-kinetic parameters, with particular attention to blastomeres and their cytoplasm. In accordance with the updated Istanbul Consensus (*3*), blastomere number and relative size are considered standard morphological criteria for evaluating cleavage-stage embryos. To examine these features noninvasively, we segmented individual blastomeres at their focal planes within reconstructed HT volumes and delineated cytoplasmic regions by excluding nuclei (Fig. 3A, see Methods). Representative HT images illustrate segmented blastomeres and cytoplasm in matured embryos, which developed to expanded blastocysts within 72 h, and in arrested embryos, which failed to progress (Fig. 3B). This segmentation enabled systematic and quantitative comparisons of blastomere morphology and cytoplasmic organization between the two groups.

**Fig. 3:**
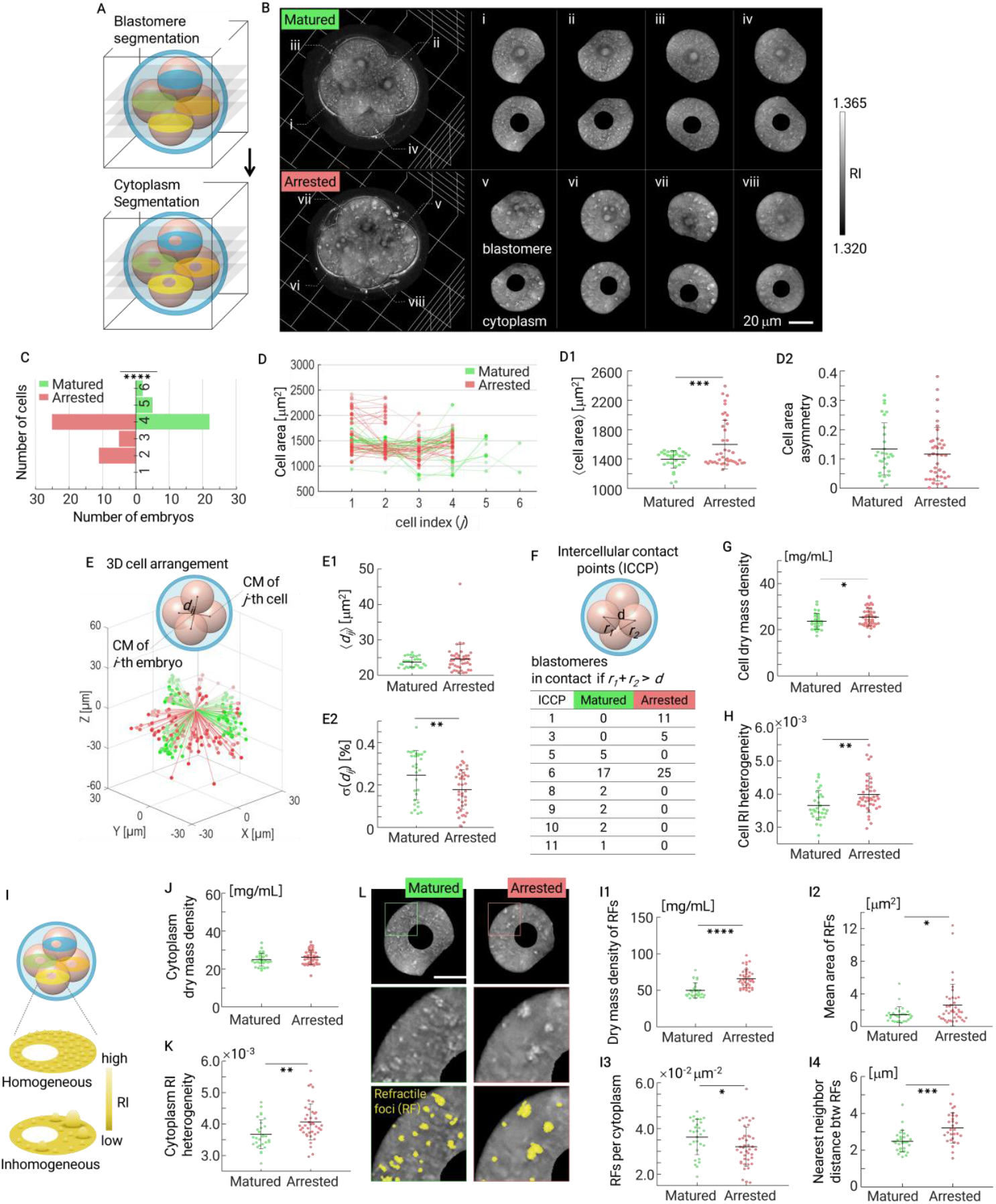
Noninvasive 3D subcellular assessment of embryo quality at 24 h post–2-cell stage. **A**, Workflow for 3D segmentation of blastomeres (top) and cytoplasm (bottom) from HT images. **B**, Representative segmented blastomeres and cytoplasm of matured (green) and arrested (red) embryos. Panels (i–iv) show blastomeres of a matured embryo, while panels (v–viii) show blastomeres of an arrested embryo. **C**, Number of blastomeres per embryo, showing reduced blastomere counts in arrested embryos. **D**, Blastomere size at focal planes: mean (**D1**) and asymmetry (coefficient of variation, CV, **D2**). **E**, Schematic of 3D blastomere spatial arrangement and centroid displacement relative to the embryo center of mass (CM); mean (**E1**) and spatial variability (CV, **E2**) of blastomeres, with arrested embryos showing reduced spatial variability. **F**, Quantification of intercellular contact points (ICCPs) per embryo, with arrested embryos more frequently exhibiting fewer contacts. **G–H**, Blastomere dry mass density (G) and RI heterogeneity (H), revealing higher RI heterogeneity in arrested embryos. **I**, Schematic illustrating cytoplasmic RI distribution: homogeneous versus inhomogeneous patterns. **J**, Comparison of mean cytoplasmic dry mass density between matured and arrested embryos, showing no significant difference. **K**, Cytoplasmic RI heterogeneity (coefficient of variation) is significantly elevated in arrested embryos. **L**, Representative HT images of cytoplasmic segmentation in the matured and arrested embryos, with refractile foci (RF) highlighted in yellow. **I1–I4**, Quantitative characterization of RFs: dry mass density (**I1**), mean area (**I2**), number per cytoplasmic area (**I3**), and nearest-neighbor distance (**I4**). Arrested embryos displayed higher dry mass density, larger foci, reduced density, and greater spacing of RFs compared to matured embryos. Data represent four independent experiments (29 matured and 41 arrested embryos). Statistical significance was assessed using unpaired or Welch’s t-tests, depending on variance equality. \**p* < 0.05, \*\**p* < 0.01, \*\*\**p* < 0.001.

Quantitative analysis of blastomere number revealed that matured embryos typically reached the 4-cell stage by 24 h, whereas arrested embryos often lagged with fewer cells (Fig. 3C). Consistent with reduced cleavage, blastomeres in arrested embryos were significantly larger in area (1,595 ± 330 μm^2^ in matured vs. 1,395 ± 123 μm^2^ in arrested; Fig. 3D1), reflecting fewer cells sharing the same cytoplasmic volume. We next assessed size asymmetry by calculating the coefficient of variation across blastomeres, which did not differ significantly between groups (0.134 ± 0.088 in matured vs. 0.116 ± 0.092 in arrested; Fig. 3D2). Although unequal blastomere size is traditionally regarded as a negative morphological feature, our analysis suggests that 3D blastomere counts provide a more robust indicator of developmental potential than size variability alone.

We next examined the 3D spatial arrangement of blastomeres by calculating the displacement of each blastomere centroid relative to the overall embryonic centroid (Fig. 3E, see Methods). The mean displacement did not differ significantly between groups (Matured: 23.69 ± 1.43 μm; Arrested: 24.58 ± 4.08 μm; Fig. 3E1). However, the coefficient of variation, representing spatial variability, was higher in matured embryos (0.246 ± 0.116) compared to arrested embryos (0.178 ± 0.093; Fig. 3E2). These results suggest that while the overall distances of blastomeres from the embryo centroid were similar, matured embryos exhibited greater spatial variability, whereas arrested embryos showed a more rigid and constrained blastomere arrangement. This finding indicates that spatial organization during early cleavage may influence subsequent developmental potential (*50, 51*).

Consistent with this interpretation, analysis of intercellular contact points (ICCPs) revealed that arrested embryos frequently had fewer contacts among blastomeres (Fig. 3F). Two blastomeres were considered in contact if the distance between their centroids was less than the sum of their radii (derived from *A*_*cell*_ = *πr*^*2*^). Arrested embryos more often exhibited fewer than six ICCPs, indicating reduced opportunities for cell–cell interaction (*52*). These findings suggest that limited intercellular contacts— potentially reflecting lower blastomere counts or weaker cell-cell interactions—may be associated with reduced developmental potential.

Beyond cellular morphology, HT enabled quantitative assessment of subcellular mass distributions. Because RI is directly proportional to dry mass density (*53*), we analyzed both mean blastomere density and RI heterogeneity at 24 h (Fig. 3G–H, see Methods). The mean dry mass density was slightly higher in arrested embryos (23.65 ± 3.46 mg/mL in matured vs. 25.48 ± 3.92 mg/mL in arrested), while the coefficient of variation—representing RI heterogeneity—showed a more pronounced difference (3.66 ± 0.45 × 10^-3^ in matured vs. 3.99 ± 0.55 × 10^-3^ in arrested). The elevated heterogeneity in arrested embryos suggests cytoplasmic irregularities that are not detectable with conventional bright-field microscopy, prompting a more detailed investigation of cytoplasmic organization.

We next examined cytoplasmic RI distribution, where arrested embryos displayed more pronounced non-uniformity (Fig. 3I). Although mean cytoplasmic dry mass density did not differ significantly between groups (Matured: 24.86 ± 3.44 mg/mL; Arrested: 26.08 ± 3.76 mg/mL; Fig. 3J), RI heterogeneity was markedly higher in arrested embryos (Matured: 3.67 ± 0.46 × 10^−3^; Arrested: 4.06 ± 0.58 × 10^−3^; Fig. 3K). To further characterize these differences, we segmented refractile foci (RFs), defined as high-RI substructures with clear boundaries (see Methods, Fig. 3L). Arrested embryos exhibited RFs with greater dry mass density (49.78 ± 10.18 vs. 65.72 ± 12.40 mg/mL; Fig. 3I1) and larger size (1.43 ± 0.97 vs. 2.59 ± 2.58 μm^2^; Fig. 3I2), but fewer foci per cytoplasmic area (3.60 ± 0.77 × 10^-3^ vs. 3.18 ± 0.86 × 10^-3^ μm^-2^; Fig. 3I3) and greater nearest-neighbor distances (2.48 ± 0.58 vs. 3.21 ± 0.82 μm; Fig. 3I4). Collectively, these findings indicate that the elevated cytoplasmic RI heterogeneity in arrested embryos arises from a sparse and irregular distribution of high-density RFs, in contrast to the more uniform arrangement observed in matured embryos.

Collectively, these findings demonstrate that HT enables quantitative, noninvasive analysis of both cellular and subcellular features during the cleavage stage. Arrested embryos were distinguished by delayed cleavage, fewer intercellular contacts, and elevated cytoplasmic heterogeneity—abnormalities likely to impair compaction and subsequent blastocyst formation. By capturing such multidimensional parameters in three dimensions, HT offers a more comprehensive evaluation of embryo quality than conventional morphology, highlighting its potential as a powerful tool for embryo selection.

### Nuclear characterization at 36 h post–2-cell stage

To extend our analysis beyond cleavage-stage embryos, we examined nuclear organization at 36 h post–2-cell stage, a period when embryos were typically entering the early morula stage. This developmental phase is often underrepresented in IVF studies due to limited imaging clarity. Using the present method, we were able to clearly resolve individual nuclei in embryos at this stage (Fig. 4). Volumetric nuclear maps were reconstructed from multiple axial RI slices by segmenting across focal planes (*Δz* = 0.88 μm), and the resulting masks were merged along the axial axis to generate complete 3D representations (see Methods). Representative 3D renderings of matured and arrested embryos at 36 h are shown in Fig. 4B, with segmented nuclei pseudo-colored in yellow for clarity. Complementary *xy*-slices further highlight the spatial arrangement of nucleoli within individual nuclei. This approach enabled precise visualization of nuclear boundaries in both matured and arrested embryos, providing a robust framework for quantitative comparisons.

**Fig. 4:**
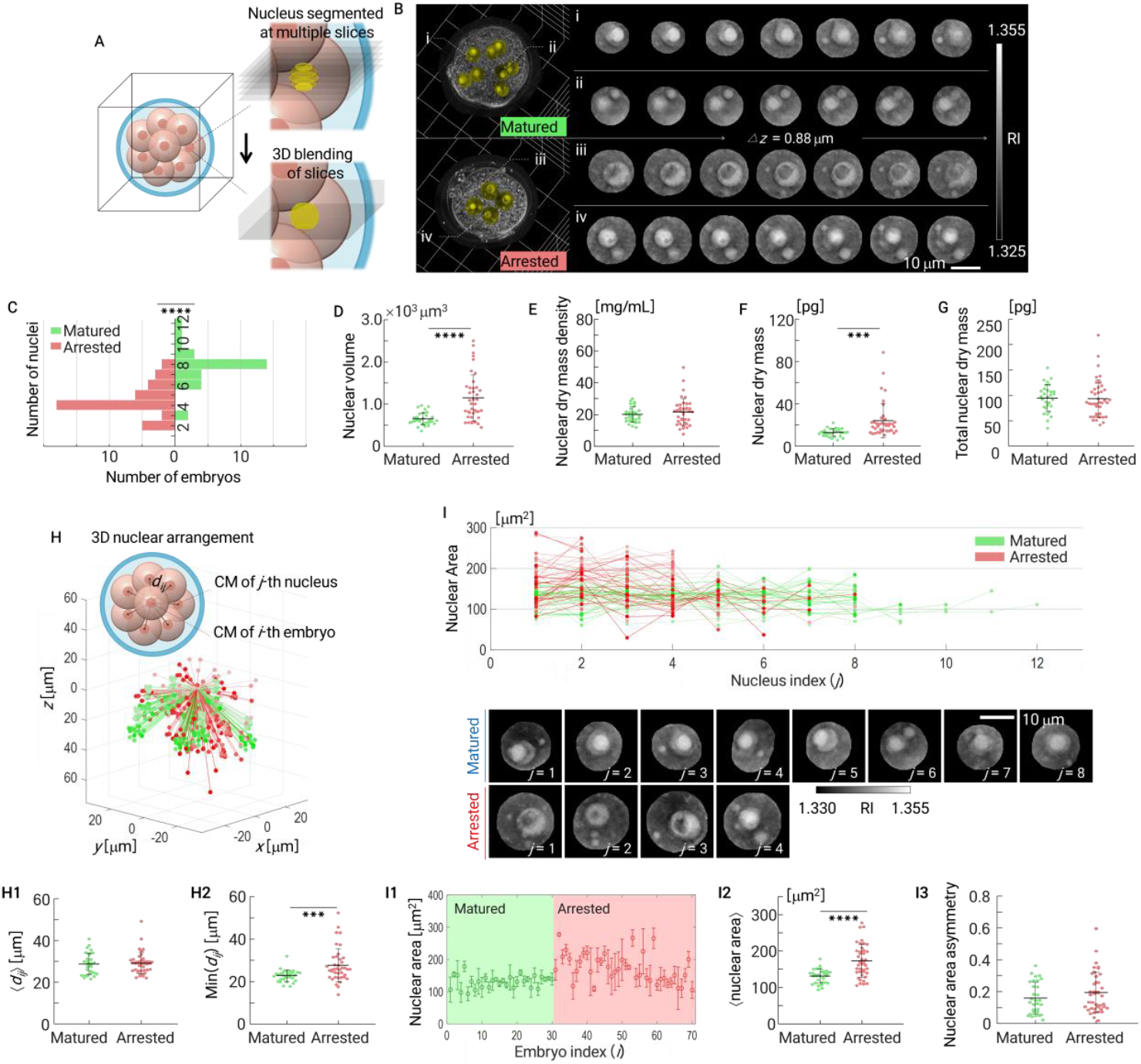
Noninvasive 3D subcellular assessment of embryo quality at 36 h post–2-cell stage. **A**, Workflow for nuclear segmentation from multiple axial RI slices, enabling volumetric reconstruction of nuclei. **B**, Representative HT images and segmented nuclei (yellow) in matured and arrested embryos. Panels (i–iv) show nuclear segmentation across focal planes (Δ*z* = 0.88 μm). **C**, Number of nuclei per embryo, showing significantly fewer nuclei in arrested embryos. **D–F**, Quantitative analysis of nuclear properties per embryo: mean values of volume (**D**), dry mass density (**E**), and dry mass (**F**). **G**, Total nuclear dry mass per embryo, which remained comparable between groups. **H**, 3D spatial displacement of nuclei relative to the embryonic centroid, showing schematic (left) and scatter plots of displacement distances. **H1**, Mean displacement of nuclei from the embryonic centroid, showing no significant difference between groups. **H2**, Minimum nearest-neighbor distance between nuclei, significantly increased in arrested embryos, indicating looser nuclear packing. **I**, Nuclear area measurements of individual nuclei (index *j*) in matured and arrested embryos, showing consistently larger nuclear areas in arrested embryos. Representative RI tomograms of nuclei from matured and arrested embryos are shown below, with indices *j*. **I1**, Nuclear area distribution across all embryos, grouped by developmental outcome. **I2**, Mean nuclear area per embryo (index *i*), significantly larger in arrested embryos. **I3**, CV of nuclear area per embryo, showing no significant difference between groups. Data were obtained from four independent experiments (30 matured and 40 arrested embryos). Statistical significance was determined by unpaired or Welch’s t-tests depending on variance equality. \**p* < 0.05, \*\**p* < 0.01, \*\*\**p* < 0.001, \*\*\*\**p* < 0.0001.

Quantification of nuclear number revealed a clear distinction between groups. Matured embryos contained significantly more nuclei, consistent with more advanced progression within the same time window (Fig. 4C). In contrast, arrested embryos frequently failed to reach the expected nuclear count and exhibited larger nuclear volumes (Matured: 651 ± 149 μm^3^; Arrested: 1,145 ± 546 μm^3^; Fig. 4D), reflecting delayed or incomplete divisions. These quantitative differences in nuclear count and volume highlight divergent nuclear organization indicative of altered cleavage and compaction states between the two groups.

We next analyzed nuclear properties at the individual cell level (Figs. 4D–G, see Methods). Nuclear dry mass density did not differ significantly between groups (Matured: 20.04 ± 4.85 mg/mL; Arrested: 21.54 ± 8.41 mg/mL; Fig. 4E). However, when nuclear volume was considered, arrested embryos exhibited significantly greater nuclear volume and nuclear dry mass per nucleus (Matured: 12.74 ± 3.59 pg; Arrested: 23.91 ± 16.41 pg; Fig. 4D, F). Despite these differences, total nuclear dry mass per embryo remained comparable (Matured: 94.57 ± 26.74 pg; Arrested: 93.24 ± 37.25 pg; Fig. 4G), indicating that overall nuclear biomass was conserved but distributed into fewer, larger nuclei in arrested embryos. These findings suggest that while impaired cleavage divisions disrupt the balance between nuclear size and number, the overall nuclear mass remains conserved, consistent with the reductive nature of early mammalian cleavage (*54, 55*). The spatial arrangement of nuclei further distinguished matured from arrested embryos (Fig. 4H).

Using the same centroid-based displacement analysis applied to blastomeres (see Fig. 3 and Methods), we measured the 3D distance of each nucleus from the embryonic centroid. The mean displacement did not differ significantly between groups (Matured: 28.72 ± 5.20 μm; Arrested: 29.17 ± 5.17 μm; Fig. 4H1). However, the minimum nearest-neighbor distance between nuclei was significantly greater in arrested embryos (Matured: 22.80 ± 2.77 μm; Arrested: 27.63 ± 7.94 μm; Fig. 4H2), indicating looser nuclear packing. These results suggest that although nuclei in both groups occupy a similar radial range, the reduced number of nuclei in arrested embryos leads to less compact organization.

Analysis of nuclear areas provided further insight into differences between matured and arrested embryos at the morula stage (Fig. 4I). Nuclear area measurements across individual nuclei (index j) consistently showed larger values in arrested embryos compared with matured embryos. Representative RI tomograms confirmed this pattern, illustrating smaller, more compact nuclei in matured embryos and larger, irregular nuclei in arrested embryos. Quantitative analysis revealed that the mean projected nuclear area was significantly larger in arrested embryos (Matured: 130.8 ± 19.5 μm^2^; Arrested: 172.7 ± 45.3 μm^2^), while the coefficient of variation of nuclear area was comparable between groups (Matured: 0.159 ± 0.099; Arrested: 0.194 ± 0.127). Aggregated distributions across embryos further emphasized this distinction (Fig. 4I1), with arrested embryos exhibiting significantly greater mean nuclear areas (Fig. 4I2) but no notable difference in size asymmetry (Fig. 4I3). Together, these results indicate that arrested embryos are characterized by fewer but disproportionately larger nuclei, consistent with altered nuclear organization arising from disrupted cleavage progression and aberrant compaction behavior.

### Leveraging early-stage 3D subcellular features for blastocyst prediction

To evaluate whether quantitative HT-derived parameters could predict developmental potential, we implemented a machine learning framework based on features extracted at 24 h and 36 h (Fig. 5A–B). HT imaging enabled systematic quantification of cellular and subcellular properties at these time points, generating a multidimensional dataset for predictive modeling. We developed a predictive model for blastocyst formation that leverages 3D subcellular features obtained during early development. The workflow comprised four steps: (i) feature extraction, (ii) feature selection, (iii) classifier construction, and (iv) blastocyst prediction (Fig. 5A). While the overall framework resembles conventional feature-based machine learning approaches, our model uniquely incorporates 3D subcellular features derived from quantitative RI analysis. Unlike traditional methods that rely primarily on 2D annotations or kinetic parameters, HT provides direct volumetric measurements of subcellular structures—including cytoplasmic dry mass density, RI heterogeneity, nuclear volume, and nuclear dry mass—enabling more comprehensive embryo characterization.

**Fig. 5:**
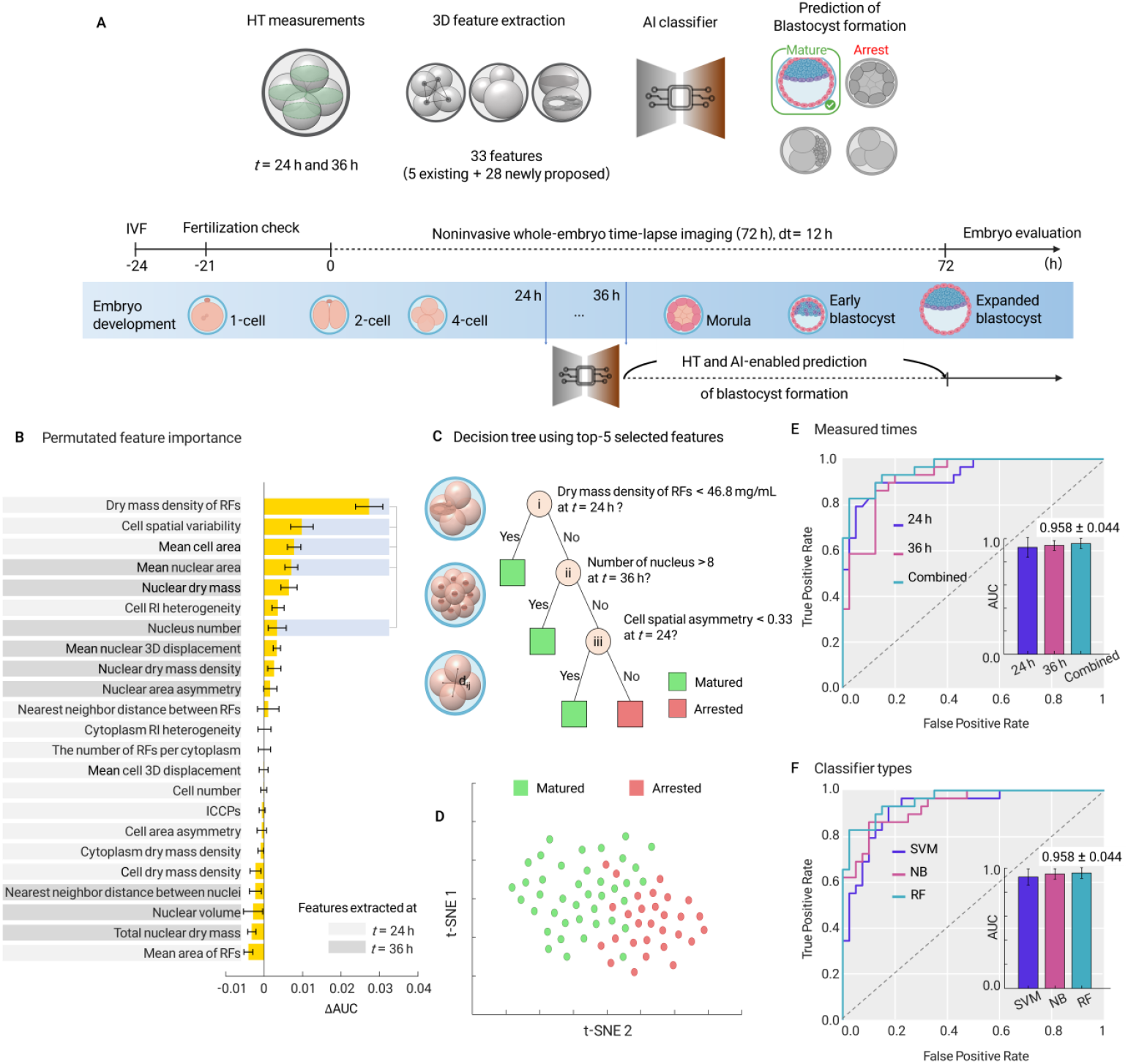
Predictive modeling of blastocyst formation using 3D subcellular features. **A**, Schematic workflow illustrating embryo development from the 1-cell to blastocyst stage (bottom) and the analysis pipeline using HT imaging performed at 24 h and 36 h for feature extraction and machine learning–based prediction of developmental potential (top). **B**, Permutated feature importance analysis from random forest classifiers, highlighting the most informative 3D features at 24 h and 36 h, including cell displacement variability, nucleus number, mean nuclear area, and refractile foci characteristics. **C**, Decision tree generated using the five most stable predictive features: (i) coefficient of variation (CV) of cell 3D displacement, (ii) nucleus number, (iii) mean cell area, (iv) CV of nuclear displacement, and (v) dry mass density of refractile foci. **D**, t-SNE visualization of embryo classification based on combined 3D features, showing separation between matured and arrested embryos. **E**, ROC curves and AUC values for prediction models trained with features from 24 h, 36 h, and combined datasets. Combined features yielded the highest accuracy (AUC [95% CI] = 0.958 [0.914–1.00]). **F**, ROC curves, and AUC values for different classifiers—support vector machine (SVM), naïve Bayes (NB), and random forest (RF) —all showing high predictive performance, with RF achieving the best accuracy.

Feature importance analysis using random forest classifiers revealed that multiple parameters contributed substantially to prediction accuracy (Fig. 5C), including fourteen features at 24 h, nine features at 36 h, and a combined set of twenty-three features. At 24 h, cell displacement variability, blastomere morphology, and cytoplasmic heterogeneity were the most informative, whereas at 36 h, nuclear number and nuclear area provided the strongest discrimination. Refractile foci properties, such as dry mass density, also ranked highly among predictive features. Together, these results indicate that both cellular organization and cytoplasmic substructure provide complementary information for assessing embryo developmental potential.

To establish an interpretable framework, we constructed a decision tree model based on the five most stable features: coefficient of variation of cell displacement, nucleus number, mean cell area, coefficient of variation of nuclear displacement, and refractile foci dry mass density (Fig. 5D). This model effectively distinguished matured from arrested embryos, supporting the notion that a relatively small subset of biologically meaningful features can capture the key determinants of blastocyst formation.

Visualization of the feature space using t-SNE demonstrated clear separation between matured and arrested embryos when features from 24 h and 36 h were combined (Fig. 5E). Prediction performance was evaluated using random forest classifiers with 10-fold cross-validation, and Receiver operating characteristic (ROC) curves were generated from pooled predictions (Fig. 5F, see Methods). Three feature sets were compared: fourteen features at 24 h, nine features at 36 h, and the combined set of twenty-three features. The mean area under the curve (AUC) values with 95% confidence intervals (CIs) were 0.925 (95% CI, 0.839–1.00), 0.942 (95% CI, 0.899–0.985), and 0.958 (95% CI, 0.914–1.00) for 24 h, 36 h, and combined features, respectively. These results underscore the added value of integrating multiple developmental stages, with the combined feature set achieving the highest accuracy.

Finally, we benchmarked the performance of different classifiers. Random forest (*56*), support vector machine (SVM) (*57*), and naïve Bayes (NB) (*58*) all achieved high predictive accuracy, with random forest consistently performing best (AUC [95% CI] = 0.958 [0.914–1.00]; Fig. 5G, see Methods). Using the combined feature set, prediction performance was similarly robust across classifiers (AUC [95% CI]: SVM, 0.925 [0.859–0.991]; NB, 0.950 [0.906–0.994]; RF, 0.958 [0.914–1.00]), again with random forest yielding the highest accuracy. Together, these findings demonstrate that HT-derived 3D subcellular features, particularly when integrated across multiple early developmental stages, enable robust and interpretable prediction of blastocyst formation. This proof-of-concept framework underscores the potential of quantitative HT and AI to augment embryo selection in IVF practice.

## Discussion

This study demonstrates that HT enables quantitative, label-free, 3D assessment of preimplantation mouse embryos and that early subcellular features are informative for predicting blastocyst formation. Across 72 h time-lapse imaging, HT resolved canonical milestones—from cleavage and compaction to blastocoel expansion—while preserving developmental competence within MEA safety criteria. At 24 h, matured and arrested embryos diverged most clearly: matured embryos showed higher blastomere counts, greater spatial variability in blastomere positioning, and more intercellular contacts; arrested embryos exhibited enlarged blastomeres and elevated RI heterogeneity within the cytoplasm. At 36 h, matured embryos presented more numerous, smaller, and tightly packed nuclei, whereas arrested embryos had fewer but larger nuclei with increased nearest-neighbor distances, indicating looser packing and reduced coordination of cellular organization.

A central biological insight is that cytoplasmic organization—not merely gross cell count or symmetry—differs systematically with outcome. Arrested embryos displayed higher cytoplasmic RI heterogeneity driven by sparser, larger, and denser refractile foci, while mean cytoplasmic dry-mass density remained similar between groups. These RF phenotypes may reflect perturbed subcellular remodeling, potentially involving changes in lipid handling, organelle clustering, or phase separation, which could influence metabolic efficiency during early cleavage. In parallel, nuclear metrics at 36 h indicate that total nuclear biomass per embryo is conserved but partitioned differently: matured embryos distribute mass across more, smaller nuclei, whereas arrested embryos retain mass in fewer, larger nuclei, with looser nuclear packing indicative of aberrant compaction.

Beyond description, we provide an interpretable predictive framework for blastocyst formation that integrates multi-scale 3D features from 24 h and 36 h. Random-forest models achieved high performance (AUC up to 0.958), and t-SNE visualizations showed clear separation between outcomes when combining stages. Importantly, a compact decision tree built from five stable, biologically meaningful features—cell-displacement variability (24 h), mean cell area (24 h), RF dry-mass density (24 h), nucleus number (36 h), and mean nuclear area (36 h)—retained strong discriminative power. This balance between accuracy and interpretability is critical for clinical translation, supporting explainable AI that can be audited against embryological knowledge rather than opaque end-to-end predictors.

These results suggest a practical translational path in IVF: HT could operate alongside time-lapse bright-field systems to add orthogonal 3D subcellular information (cytoplasmic heterogeneity, nuclear packing) that is invisible to 2D imaging, thereby refining selection beyond morphokinetics alone. Because HT is label-free and met safety benchmarks in mouse embryo assays, it is well-positioned for preclinical validation in human embryos under research protocols. In the near term, the most feasible integration is periodic HT snapshots at key inflection points (e.g., ∼24 h and ∼36 h) to minimize dose while maximizing information gain, with automated pipelines extracting the small, validated feature set required by the decision tree.

Integrating dynamic aspects of live embryo imaging would further expand the repertoire of developmental indicators, such as cleavage timing, cell-cycle duration, and division synchrony. In this study, however, the imaging interval for time-lapse acquisition was set to 12 h to minimize total light exposure, even though HT can capture full 3D images every 7 s, enabling continuous monitoring of embryogenesis. This conservative interval was chosen because the imaging field of view is smaller than the illumination area, necessitating tile imaging in group culture and thereby increasing total exposure (see Methods). By contrast, single-embryo culture could reduce exposure and allow shorter imaging intervals under the same light dose. Additional reductions in light exposure may be achieved by lowering illumination intensity per slice—facilitated by cameras with higher signal-to-noise ratios—or by decreasing the number of slices required to reconstruct each 3D RI tomogram. The latter could be realized through advanced strategies such as optimized illumination schemes (*59*) or spectral multiplexing (*60*), which would enhance acquisition efficiency while maintaining reconstruction fidelity.

We also recognize limitations. First, the dataset—though multi-experiment—is modest, raising concerns of overfitting despite stratified cross-validation; independent cohort and leave-one-experiment-out validations are warranted, ideally across mouse strains and laboratories. Second, our time-lapse cadence (12 h) sacrifices the temporal resolution of morphokinetic data to limit exposure during tiled group-culture imaging; single-embryo culture, improved camera SNR, optimized illumination, or slice-count reduction (e.g., multiplexed transfer-function reshaping) could enable shorter intervals under equal or lower dose. Third, RFs are defined operationally by RI and gradients; biochemical identity (lipid droplets vs. glycogen bodies vs. organelle clusters) remains putative and should be probed via correlative assays (e.g., gentle live dyes in animal models, cryo-EM/label-free correlative modalities, or post-fix verification in research contexts).

Methodologically, several components are primed for automation. SAM-assisted and manual segmentations demonstrate feasibility but are not scalable for routine use; supervised or weakly supervised 3D segmentation models trained on HT volumes can standardize blastomere, nuclear, and RF extraction, reduce operator variability, and support prospective clinical studies. Likewise, the current feature set—while already compact—could be stress-tested with embedded controls (illumination dose, culture density) and calibrated across instruments to ensure reproducibility. Reporting standards should include absolute timing, dose budgets per stack, voxel-size calibration, and uncertainty estimates for RI-to-mass conversion (*α* variability across biomolecular composition).

In summary, HT enables quantitative 3D phenotyping that links early subcellular architecture—such as cytoplasmic heterogeneity and nuclear packing—to downstream developmental competence, and these features can be distilled into interpretable predictors of blastocyst formation. Our findings establish a foundation for fully automated, noninvasive embryo evaluation in IVF. To improve scalability, steps that currently require manual handling, such as segmentation of subcellular structures, could be automated with AI (*40, 61, 62*), while coupling with virtual staining may provide molecule-specific insights to further enhance interpretability (*61*).

Future work should (i) validate the model across independent mouse cohorts and multi-center datasets, (ii) conduct carefully governed studies in human embryos to assess generalizability, (iii) investigate the biophysical basis of refractile foci heterogeneity and nuclear-packing defects, and (iv) integrate HT with clinic-ready automation and decision support. If confirmed, HT-augmented embryo assessment could shift current practice from largely qualitative 2D grading toward objective, mechanistically grounded, and explainable 3D selection criteria. Further exploration of the mechanical and hydraulic regulation of early mammalian embryos may benefit from conceptual parallels drawn from ovarian and follicular mechanobiology (*63, 64*).

Our study establishes HT as a powerful platform that integrates whole-embryo volumetric imaging with subcellular biophysical profiling, offering a complementary and objective framework to enhance traditional morphology-based embryo assessment. By linking early subcellular architecture to developmental competence and distilling these signatures into interpretable predictors, we demonstrate the feasibility of objective embryo assessment that extends beyond conventional morphology. With continued advances in AI-driven analysis and virtual staining, HT has the potential to transform IVF practice, enabling embryo selection that is not only explainable and mechanistically anchored but also clinically actionable.

## Materials and Methods

### Embryo culture

All mice used for conventional IVF were purchased from Orient Bio Inc. (Sungnam, South Korea). For the low-coherence HT embryo assay, 4-week-old F1 hybrid females (C57BL/6 × DBA/2) and 10-week-old F1 hybrid males (C57BL/6 × C3H) were employed, whereas 4-week-old BALB/c females and 8-week-old BALB/c males were used for noninvasive imaging analysis. In each experiment, seven females and two males were utilized. Animals were maintained on standard rodent chow with ad libitum access to reverse osmosis water. All experimental procedures were approved by the Institutional Animal Care and Use Committee (IACUC) of KAIST (KA2023-005-v4).

Superovulation was induced by intraperitoneal injection of 0.1 mL CARD Hyperova (Cosmo Bio, Japan), followed 48 h later by 20 IU human chorionic gonadotropin (hCG; Sigma-Aldrich, USA). At 15–18 h post-hCG, females were sacrificed by cervical dislocation and oviducts excised. Males were euthanized by cervical dislocation and sperm collected from the caudal epididymis. Cumulus–oocyte complexes were isolated from oviducts and cultured in 100 μL drops of human tubal fluid (HTF; Cosmo Bio, Japan) overlaid with 10 mL mineral oil (GeorgiaChem, Korea). Sperm were released into 100 μL drops of preincubation medium (PM; Cosmo Bio, Japan) overlaid with mineral oil and preincubated for 30 min. Fertilization was performed by adding 3 μL of sperm suspension into HTF drops containing oocytes.

At 3 h post-insemination, zygotes were collected, washed three times in HTF, and cultured overnight (∼21 h). Two-cell embryos were then transferred into Oosafe 4-well dishes (SparMed, Denmark), with 20– 30 embryos per 20 μL drop of KSOM medium (Cosmo Bio, Japan) overlaid with 0.8 mL mineral oil per well. For each experiment, two independent dishes were prepared: one for HT imaging and one for control culture. Dishes were maintained either in the incubation chamber of the low-coherence HT system (Tokai Hit, Japan) or in a conventional incubator, both set to 37°C with 5% CO_2_ under humidified conditions.

### Optical setup of low-coherence HT

To monitor mouse preimplantation development, we employed a customized HT system (HT-X1, Tomocube) optimized for embryo imaging (see Fig. S1 and Supplementary Materials). The illumination source was a light-emitting diode (LED; LE A Q8WP, OSRAM) with a peak wavelength of 624 nm, previously reported to be safe for embryo assessment (*65*). The collimated beam was modulated in the Fourier plane by a digital micromirror device (DMD; DLP4500, Texas Instruments) and relayed through a lens (CPL150, Tomocube) to project patterned illumination onto the back focal plane of a condenser lens (NA 0.72; CWD30, Tomocube). Four optimized illumination patterns were combined with axial scanning, and the resulting intensity images were reconstructed into 3D RI tomograms via deconvolution (*18, 66*).

The patterned light was projected onto the sample, with an average intensity of 13.37 W/m^2^ measured using an optical power meter (PM100D, Thorlabs). During time-lapse imaging, embryos received a cumulative light dose of 1.8 kJ/m^2^, comparable to reported safe ranges for red illumination. Transmitted light was captured by a CMOS camera (FS-U3-28S5, FLIR) using an objective lens (NA 0.7; UCPLFLN20X, Olympus) and a tube lens (AC508-180-A, Thorlabs). Each RI tomogram encompassed a volume of 238 × 238 × 125 μm^3^, reconstructed from four intensity images per axial slice across 120 z-positions. The final RI tomograms achieved lateral and axial resolutions of 176 nm and 1.09 μm, respectively (*67*).

### Image acquisition and annotation

To capture the entire region containing group-cultured embryos, wide-field RI images (1.5 × 1.5 mm^2^ field of view) were obtained by stitching 7 × 7 tiles. Time-lapse imaging was performed for 72 h at 12-h intervals, covering development from the 2-cell stage to the expanded blastocyst in both brightfield and HT modes. For each experiment, embryos cultured in the same well of an Oosafe 4-well dish were imaged. In total, 85 time-lapse HT datasets were acquired across four independent experiments.

Morphological annotation was performed by experienced embryologists. For each embryo, maximum-intensity projection (MIP) images and six representative XY slices of the 3D RI tomograms were reviewed. Based on these evaluations, embryos were classified into four categories: (i) 30 embryos that progressed to the expanded blastocyst within 72 h (Matured), (ii) 11 embryos that reached the early blastocyst stage, (iii) 41 embryos that failed to reach the blastocyst stage (Arrested), and (iv) 3 embryos excluded due to loss of focus at the endpoint. For noninvasive analysis, only the Matured and Arrested groups were included. At selected developmental time points (*t* = 24 h and *t* = 36 h), 1–2 HT images were further excluded when acquisition errors occurred, such as severe embryo displacement during tile scanning.

### Image segmentation

For 3D segmentation of blastomeres and their subcellular structures, we used the Segment Anything Model (SAM; https://segment-anything.com/), an open-source framework. For each embryo, 2D masks were generated at 15–20 axial planes using SAM in combination with manual correction. SAM provided accurate segmentation at in-focus planes, whereas manual annotation was required at out-of-focus planes where embryo boundaries were indistinct. Segmented XY slices were then blended along the axial direction using the open-source Blended 3D poly2mask function (MATLAB Central File Exchange, 2013), producing continuous 3D masks.

For noninvasive analysis at *t* = 24 h, blastomeres were segmented at their focal planes using SAM with manual refinement. Cytoplasmic regions were delineated by subtracting nuclear areas, and refractile foci were identified using a rule-based approach. The cytoplasm was divided into two complementary regions: a dense interior (core) obtained by morphological erosion, and a peripheral boundary defined as the remaining cytoplasmic area. To account for edge-related noise, different RI gradient thresholds were applied (40% of the maximum local RI gradient for the core and 50% for the boundary). Core- and boundary-derived masks were then combined, and an RI threshold (>1.340) was applied to retain only high-RI refractile foci. Morphological refinement was performed to smooth boundaries, and objects smaller than 0.5 μm^2^ were excluded. All segmentation of refractile foci was implemented in MATLAB R2025a. For noninvasive analysis at *t* = 36 h, nuclei were segmented manually. Each nucleus was annotated across 3–7 axial planes, and the *xy*-slices were smoothed along the axial axis to construct continuous 3D nuclear masks.

### Quantitative image analysis

All analyses were performed in MATLAB R2025a. Dry mass was estimated from the linear relation between the RI and dry mass density, *n*(**r**) = *n*_*m*_ + *α*C(**r**), where *n*(**r**) is the RI of the sample, *n*_*m*_ is the RI of the KSOM medium (approximated to 1.336), *α* is the RI increment, and C(**r**) is the dry mass density. Dry mass density was derived from the mean RI of blastomeres and their subcellular structures using *α* = 0.18 ml g^-1^, a commonly accepted value for aqueous biomolecular samples. Within the nuclear masks, RI values below that of the medium were set to the medium RI to prevent nonphysical negative estimates. Dry mass was then calculated by multiplying the summed RI values of the segmented region by its physical volume, determined from the voxel number and the unit voxel size.

The 3D spatial organization of blastomeres and their subcellular structures within embryos was analyzed as follows. To calculate the centroid of each embryo, we first identified the XY slice with the maximum cross-sectional area from the segmented 3D masks and computed its center of mass. This approach was adopted to minimize bias from out-of-focus XY slices and to prevent over-weighting regions with larger segmented volumes, thereby providing a more robust estimate of embryo position. The center of mass of individual blastomeres and their subcellular structures were also calculated within their respective segmented masks. Nearest-neighbor distances were computed as the minimum Euclidean distance between object centroids after excluding self-distances. For each embryo, the mean nearest-neighbor distance was calculated.

### Machine-learning analysis

We evaluated three standard machine learning classifiers: support vector machine (SVM), naïve Bayes, and random forest. Models were evaluated using stratified 10-fold cross-validation with a fixed random seed. Receiver operating characteristic (ROC) curves were generated from pooled out-of-fold test predictions, providing smooth estimates for the relatively small dataset (29 matured and 40 arrested embryos) and ensuring no information leakage across folds. Mean area under the curve (AUC) values and 95% confidence intervals (CI; calculated as 1.96 × standard error of the mean, SEM) were calculated from fold-wise results. All analyses were implemented in MATLAB R2025a using TreeBagger with 100 trees (random forest), fitcsvm (SVM with RBF kernel, automatic scaling and standardization), fitcnb (naïve Bayes), and perfcurve (ROC/AUC computation).

To identify biologically informative features, we applied a two-step feature selection pipeline to the best-performing model (random forest trained on combined features at 24 h and 36 h). First, out-of-bag permutation importance (100 trees, 10 permutations per feature) was computed, and the top eight candidate features were retained. Second, cross-validated permutation importance (10-fold, 10 permutations per feature per fold) was calculated within this candidate set. Features were ranked by frequency of appearance in the top five across folds, average rank, and mean ΔAUC, yielding the five most stable predictors. A single interpretable decision tree was then trained on these five features using fitctree (MATLAB R2025a), with complexity constrained by minimum leaf size (MinLeafSize ≈ *N*/60, where *N* is the number of training samples) and maximum number of splits (MaxNumSplits = 8).

### Statistical analysis

Statistical analyses were performed using custom scripts written in MATLAB 2025a. Data are presented as mean ± SEM. For comparisons between two groups (i.e., matured versus arrested embryos), unpaired two-tailed *t*-tests were conducted. Equality of variances was assessed using Levene’s test implemented with the vartestn function (TestType, ‘LeveneAbsolute’). When equal variances were assumed (*p* > 0.05), Student’s t-test was applied; otherwise, Welch’s t-test was used. Statistical analyses were performed at the 24-hour post–2-cell stage and the 36-hour post–2-cell stage, when developmental trajectories between groups became distinct, allowing for physiologically meaningful comparisons of 3D subcellular parameters. The number of independent experiments (*N*) and the total sample size are indicated in the corresponding figure legends. Statistical significance was defined as *p* < 0.05 (\**p* < 0.05, \*\**p* < 0.01, \*\*\**p* < 0.001, \*\*\*\**p* < 0.0001). All tests were two-sided.

## Funding

National Research Foundation of Korea (RS-2024-00442348)

Korea Institute for Advancement of Technology (KIAT) through the International Cooperative R&D program (P0028463)

Korean Fund for Regenerative Medicine (KFRM) grant funded by the Korea government (the Ministry of Science and ICT and the Ministry of Health & Welfare) (21A0101L1-12)

Korea Health Technology R&D Project through the Korea Health Industry Development Institute (KHIDI) funded by the Ministry of Health & Welfare (HR22C1605)

## Author contributions

Conceptualization: YKP, JHK. Methodology: CHL, GK, TSS, SHL, JYK, KHC, JED, JHP, JPD. Animal experiments: JED. Investigation (data acquisition and analysis): CHL. Data annotation: TSS, JYK. Data interpretation: All authors. Funding acquisition: YKP, JHK. Supervision: YKP, JHK. Writing – original draft: CHL. Writing – review and editing: All authors.

## Competing interests

JHP, JPD, and YKP have financial interests in Tomocube, a company that commercializes HT. All other authors declare no competing interests.

## Notes

### Competing Interest Statement

G.P., J.H.P., J.P.D., and Y.K.P. have financial interests in Tomocube, a company that commercializes HT and QPI products. All other authors declare no competing interests.

### Summary of Updates

All figures and corresponding main text were revised.

